# Xdrop: targeted sequencing of long DNA molecules from low input samples using droplet sorting

**DOI:** 10.1101/409086

**Authors:** Esben Bjørn Madsen, Thomas Kvist, Ida Höijer, Adam Ameur, Marie Just Mikkelsen

**Affiliations:** Samplix, Herlev, Denmark; Science for Life Laboratory, Department of Immunology, Genetics and Pathology, Uppsala University, Sweden

## Abstract

Long-read sequencing can resolve regions of the genome that are inaccessible to short reads, and therefore such technologies are ideal for genome-gap closure, solving structural rearrangements and sequencing through repetitive elements. Here we introduce the Xdrop technology: a novel microfluidic-based system that allows for targeted enrichment of long DNA molecules starting from only a few nanograms of genomic DNA. Xdrop is based on isolation of long DNA fragments in millions of double emulsion (DE) droplets, where the DE droplets containing a target sequence of interest are fluorescently labeled and sorted using flow cytometry. The final product from the Xdrop procedure is an enriched population of long DNA molecules that can be investigated by sequencing. To demonstrate the capability of Xdrop, we performed enrichment of the human papilloma virus (HPV) 18 integrated in the genome of human HeLa cells. The enriched DNA was sequenced both on long-read (PacBio and Oxford Nanopore) and short-read (Illumina) platforms. Analysis of the sequencing reads resolved three HPV18-chr8 integrations at base pair resolution, and the captured fragments extended up to 30 kb into the human genome at the integration sites. In summary, our results show that Xdrop is an efficient enrichment technology for studying complex regions of the genome where long-range information is required.

## Introduction

Short-read next-generation sequencing (NGS) technologies are based on generation of millions of DNA sequence reads, typically 100-500 bp in length. As the sequence information is broken up in millions of short sequence fragments it is crucial that the reads are assembled correctly in order not to lose sequence information. This is a particularly challenging task in regions with repetitive sequence or with structural rearrangements^1^. Long-read single molecule sequencing technologies provided by Pacific Biosciences (PacBio) and Oxford Nanopore Technologies (ONT) are capable of reading DNA molecules in a range between tens to hundreds of kilobases, thereby practically solving the issue with assembling the sequence reads correctly, even for organisms with large genome sizes, such as humans^2,3^. However, the long-read technologies come with an increased cost per sequencing unit compared to the short-read sequencing instruments. The cost can be reduced by enriching the DNA sample for the genomic regions of interest, and thereby increasing the number of reads covering those specific regions.

At present, there is a wide selection of target enrichment methods for short-read sequencing technologies (reviewed in ^4^), but only few that are compatible with the long-read sequencing platforms. The most common approach to enrich for long DNA fragments is to perform a long-range PCR (LR-PCR) followed by sequencing of the resulting amplicons^5,6^. However, it is technically challenging to amplify regions longer than 10 kb using LR-PCR and both ends of the target fragment must be known in order to design the PCR primers. An alternative approach is to perform hybridization^7^ using short DNA-or RNA-probes that are used to pull down the target of interest. Hybridization-based methods can enrich for fragments up to 10 kb based on the information of a short probe (<200bp) but require relatively large amounts of input DNA (>500 ng). Both long-range PCR and hybridization methods also have a risk of introducing chimeric reads during amplification steps during sample or library preparation. Alternative protocols have recently been developed that uses the CRISPR/Cas9 system and does not require any PCR amplification step^8,9^. The amplification-free enrichment makes it possible to interrogate genomic regions that are difficult to amplify, such as expanded repeats^9,10^, but require high amounts of input DNA and are relatively labor intensive. Thus, there is a need for a method capable of isolating long-fragments from small amounts of input DNA where only a small piece of the sequence is known, using a standardized and fast protocol.

Here we present the Xdrop technology, a novel microfluidic-based system that allows for targeted enrichment of long DNA fragments. The Xdrop system permits the enriched DNA to be sequenced on a long-read sequencing platform, such as PacBio or ONT, or alternatively to provide long-fragment information from short-read sequencing. Only ∼150 bases of sequence information is needed to perform enrichment of DNA fragments of up to 40kb in size, and the protocol require as little as 1-2 ng of DNA as input to enrich for a specific region in a human sized genome. All amplification reactions are performed in small droplets and this procedure virtually eliminates the risk of forming chimeric molecules from a specific target. As a proof-of-principle, we applied the Xdrop technology to determine the integration sites of human papillomavirus 18 (HPV18) in the HeLa cell line.

## Results

### Production of millions of double emulsion droplets using the Xdrop system

The Xdrop technology is based on the generation of millions of double emulsion (DE) water-in-oil-in-water droplets with an inner diameter of 15 µm and outer diameter of 20 µm (see Figure 1A), i.e. in the same size-range as large eukaryotic cells. The DE droplet generation is performed using a microfluidic chip that has the capacity of producing up to 3000 droplets per second (Figure 1B). Because of their small volume and high stability, the DE droplets are ideal for the Xdrop enrichment work flow. The material required for an Xdrop enrichment assay is a genomic DNA sample, millions of DE droplets, and a set of PCR primers that uniquely amplify a small piece of DNA (100-200bp) within the target region of interest and a hydrolysis probe.

**Figure 1.**
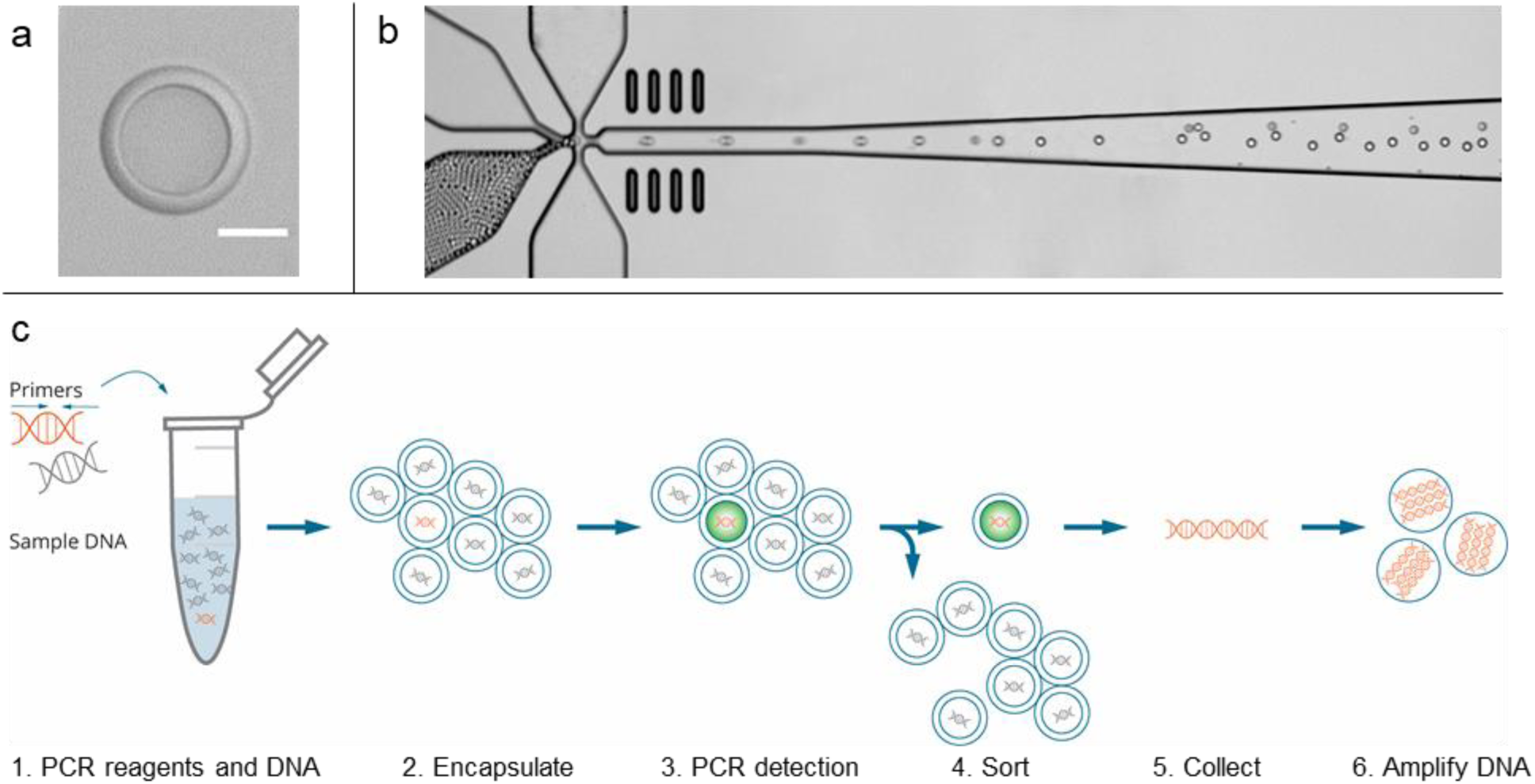
Overview of the Xdrop technology. **A**) Close-up of a double emulsion (DE) droplet. The inner droplet contains PCR-reagents and sample DNA and is surrounded by a thin oil shell. The scalebar is 10 μm. **B)** High-speed camera photo of DE droplet generation on chip. Water-in-oil droplets are being injected on the left, to generate water-in-oil-in-water droplets at the x-junction. **C)** Overview of Xdrop enrichment workflow. PCR reagents including primers are mixed with sample DNA (1) before being encapsulated in DE droplets (2). Droplet PCR allows fluorescence-based detection of the DNA molecules of interest (3) that are then sorted out on a cell sorter (4). The DNA from the sorted droplets is finally collected (5) and amplified using droplet MDA. The orange DNA helixes depicts the target DNA of interest and grey DNA helixes depicts non-target DNA.

### Isolation of DE droplets containing long DNA molecules of interest

Prior to DE droplet generation, sample DNA is mixed with PCR reagents, target specific PCR primers and a hydrolysis probe (Figure 1C). DNA and PCR-reagents are then partitioning in 34 millions of picoliter DE droplets, of which the majority will only contain either a single DNA molecule or none. The droplets are thermocycled in a standard PCR apparatus, allowing 34 million individual PCR-reactions to be performed in parallel. In the droplets containing a target DNA molecule, the target-specific primers and hydrolysis probe will bind to the template and, upon primer extension, the 5’ exonuclease activity of the polymerase cleaves the fluorophore from the probe thereby releasing a fluorescent dye. By detecting the fluorescent droplets, it is therefore possible to identify and isolate the droplets carrying the DNA molecules of interest (Figure 1C). The detection and sorting can be carried out on a standard laboratory cell sorter (FACS), as the carrier fluid of the DE droplets is aqueous and the DE droplets are sufficiently small. On the FACS apparatus, the DE droplets scatter the light and are easy to distinguished from oil-in-water droplets using forward- and side-scatter. In a second gate, the PCR positive DE droplets with high FITC/FL1 fluorescence are separated from the PCR negative droplets that are low in FITC/FL1 fluorescence. These droplets are sorted and the droplets containing the target DNA molecule of interest are collected (Figure 1C). The sorted droplets are then coalesced to release the DNA into solution. As the amount of DNA fragments collected from the sorted droplets is in the femtogram range and hence too low to be sequenced using NGS technologies, the sorted DNA fragments are amplified using Phi29 polymerase multiple displacement amplification (MDA). To avoid amplification bias and introduction of chimeric amplification products, DNA amplification of the minute amount of sorted DNA, the DNA and MDA reaction mix is emulsified, creating tens of thousands separate MDA reactions droplets. Only a fraction of the droplets will contain a DNA molecule in the first round of amplification. Therefore, the MDA emulsion is coalesced and used as template for a second round of droplet MDA (dMDA) to generate a sufficient amount of DNA.

### Xdrop enrichment for integrated HPV18 virus

To evaluate the Xdrop enrichment procedure on a human sample with biological relevance, we designed an enrichment experiment targeting human papilloma virus 18 (HPV18) integrated in the human cancer cell line HeLa. Previous studies have shown that the HeLa genome contains integrated HPV18 virus DNA^11,12^. A single primer-probe set located between HPV18 position 5916 and 6014 was designed to detect HPV18 containing DNA fragments. 1.5 ng of genomic HeLa DNA, corresponding to 417 HeLA genome copies, was partitioned into DE droplets prior to amplification with the target specific primers. The HPV18 positive DE droplets with high FITC/FL1 fluorescence were easily separated from the HPV18 negative DE droplets on the scatter plot obtained from the FACS sorting (supplementary Figure S1A). A total of 143 HPV18 positive DE droplets were sorted and the collected HPV18 DNA fragments were amplified using dMDA. After two rounds of dMDA, the total DNA content was 3 µg with an average DNA fragment length of more than 15 kb (supplementary Figure S1B). The number of HPV18 containing DNA fragments was estimated using a semi-quantitative qPCR and was found to be approximately 1000-fold higher per ng of total DNA as compared to the starting material.

### Detection of HPV18 integration sites in the HeLa genome

To generate long, continuous reads of the enriched molecules, we performed single-molecule real-time (SMRT) sequencing. A 2 kb insert library was prepared directly from the HPV18 enriched DNA sample and sequencing was performed on the PacBio RSII system. Our rationale for choosing a relatively short library size (2kb) in this experiment was to generate high accuracy circular consensus sequences (CCS)^13^ of a sufficient length to easily identify the HPV18 integration sites. The sequencing resulted in about 40,000 CCS reads, of which 306 reads (0.76%) could be mapped to the HPV18 viral genome. Without enrichment we would not expect any of the 40,000 reads to originate from HPV18, as the probability of having at least a single HPV18 containing read is below 10% (assuming the presence of 3 HPV18 integrations per genome). The HPV18 reads were covering the HPV18 gene region E1, E6-E7 and L1 but lacked the central part of the viral genome encoding E2, E4, E5 and L2 (Figure 2A and supplementary Figure S2). This is in concordance with the structure previously reported for HPV18 integrated in the HeLa genome^11^. By extracting all reads mapping to HPV18 and re-aligning them to the human reference genome, we identified four HPV18 fusion points to a region of chromosome 8 (Figure 2B). Detailed analysis revealed that one end of HPV18 is fused to three different locations in chromosome 8, while the other end of HPV18 (position 5736) always is fused to the same position on chromosome 8. The proposed structure of the HPV18 integrations in the HeLa genome is shown in Figure 2B. All of the identified HPV18-HeLa fusion points were validated by PCR and Sanger sequencing (supplementary Figure S3).

**Figure 2.**
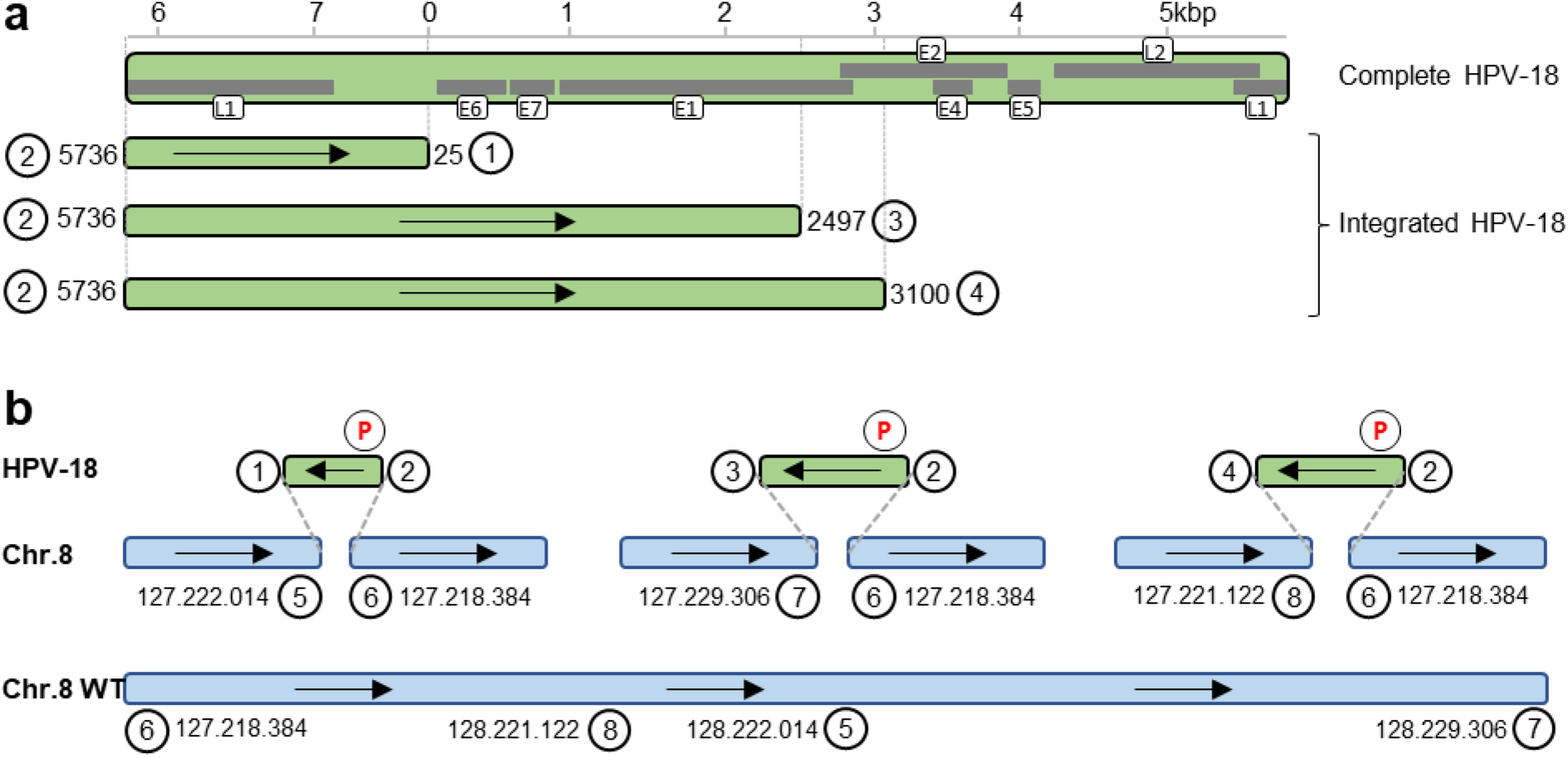
Detection of HPV18 integration sites by Xdrop enrichment and SMRT sequencing. **A)** Overview of the complete HPV18 genome with genes depicted in grey. Below the 3 types of integrated HPV18 found in the HeLa genome. The breakpoints are shown with numbered circles. **B)** Overview of the HPV18 fusion points and the suggested structure of the integrations identified in the HeLa genome. The positions of the chromosome 8 integration sites refer to the GRCh38 genome assembly. The position of the primers used for enrichment is shown with red P’s. The numbered circles correspond to the numbering in A).

To evaluate the robustness of the Xdrop method, we repeated the HPV18 enrichment from the same HeLa DNA sample and performed sequencing on Illumina and ONT instruments. The same HPV18/Chr8 fusion points were also identified using these technologies, but only three of the four fusion sites where found in the ONT data set. The coverage at each fusion point is shown in supplementary Table 1, and importantly the enriched DNA fragments extend up to 30 kb into the human genome at the integration sites (supplementary Figure S4). The percentage of on-target reads, including the reads covering the integration site region, was 5%, 7% and 10% for the PacBio, ONT and Illumina dataset, respectively. Our sequencing results demonstrate that the Xdrop system is able to enrich for very long DNA molecules, and that different types of sequencing libraries can be produced from the resulting enriched DNA.

## Discussion

The Xdrop technology is a novel targeted DNA enrichment method that isolates long DNA fragments by FACS sorting of DE droplets. Since the method makes it possible to enrich for large unknown flanking genomic regions, it is highly relevant for a wide range of applications including genome gap-closing^14^, targeted sequencing of hypervariable regions or gene families^15^, investigation of repetitive regions^16^ and as demonstrated here in identifying viral insertion sites in a genome. Also, there are several emerging clinical applications of targeted long-read sequencing^17^, where the Xdrop technology could prove to be beneficial as compared to alternative enrichment methods. The long fragments generated with the Xdrop method not only lend themselves well to long-read sequencing but can also provide valuable context information to short-read sequencing, as demonstrated by the Illumina sequencing results (supplementary Table S1).

A unique advantage of the Xdrop workflow is that the DNA amplification can be initiated from femtogram amounts of target DNA, compared to other enrichment protocols were nanograms^5,7^, or even micrograms^9,10^, of DNA is required. Performing the MDA reaction in droplets has the additional advantage that it reduces the amplification bias between different DNA fragments. Furthermore, the compartmentalization of single DNA molecules drastically reduces the risk of chimeric molecules being generated by the Phi29 polymerase.

In the current study, a FAM-labeled hydrolysis probe was used for the detection of droplets containing HPV18 DNA molecules. Multiplexing of several different DNA targets by adding additional primer- and probe-sets is expected to be feasible. The targets could then be distinguished by differently labeled probes, thereby allowing quantification of multiple targets in the same sample, which would be useful for example to measure copy number variability. Alternatively, the different targets could be labeled using the same dye or by using an intercalating dye, which might be preferable for example when tiling over a longer genomic region. It is possible to enrich for the same region in several samples and perform a multiplexed sequencing run. In that case, sample specific barcodes would need to be introduced into the library prior to sequencing.

The method described here can conveniently be applied to other targets, where structural information is sought, by design of a simple primer-set. With an efficient droplet production and optimization of the final amplification, the procedure can be completed in less than 24 hours from sample to library-ready enriched sample and will then represent an efficient and flexible alternative to hybrid capture or long-range PCR based enrichment methods.

## Acknowledgements

PacBio and Oxford Nanopore sequencing was performed by the National Genomics Infrastructure (NGI), which is hosted by SciLifeLab in Uppsala. Computations were performed on resources provided by SNIC through Uppsala Multidisciplinary Center for Advanced Computational Science (UPPMAX) under project b2017186. The work was funded by the Eurostars and EU Horizon 2020 research and innovation programme under project E!10942 DNANext.

## Methods

### PCR and droplet chemicals

HeLa DNA (New England Biolabs) was diluted with DNAase free water (Gibco) to 0.5ng per µl prior to use. PCR-mix for 20µl was set up as follows. 2µl of 10x PCR-buffer without detergent (Thermo Scientific), 2µl 25mM MgCl2 (Thermo Scientific), 2µl 2mM dNTP (Thermos scientific), 1.2µl Glycerol 50%, 0.4µl GoTaq polymerase (5U/µl) (Promega), 0.25µl (2mg/ml) Bovine Serum Albumin (Thermo Scientific), 0.8µl HPV18 fw primer (10µM) 5’- TGTGCTGGAGTGGAAATTGG-3’, 0.8µl HPV18 rev primer (10µM) 5’- GGCATGGGAACTTTCAGTGTC-3’, 0.6µl HPV18-TP1 probe (10µM) 5’- FAM- CAACACCTAAAGGCTGACCACGG-BHQ1-3’, 1ul 0.5ng/µl HeLa DNA and water to 20μl. For the primary droplet production 3% fluorosurfactant (RAN Biotechnologies) in Novec HFE-7500 was used as carrier phase. For the secondary droplets a DE-buffer containing 1.5x Optima buffer (40mM Tris-HCl, 60mM Trizma-base, 25mM (NH4)2SO4, 0.015% Tween 80, 45mM NaCl) and 3% glycerol was used as carrier phase.

### Droplet production

Double emulsion droplets were produced using a two-step emulsification procedure, initially creating water in oil (W-O) droplets followed by second emulsification to create water in oil in water (W-O-W) droplets.

### Primary droplets (W-O)

The initial chip used to prepare the primary emulsion was a 14µm etch depth hydrophobic “Small Droplet Chip, 14µm” (Dolomite Microfluidics). Liquids were pushed into the microfluidic chip using MFCS-EZ pressure controller (Fluigent, Germany) applying pressures of 640 mbar on the primary sample (PCR) and 650 mbar to the secondary liquid (Oil). Droplet production was done for approx. 40 minutes, processing a total of 40 µl PCR mixture.

### Secondary emulsions (W-O-W)

Immediately following primary production, droplets were collected in PTFE-tubes using a 1 mL syringe (Scientific Glass Engineering, Australia) ensuring an air-free liquid system to pull the droplets into the tube. The tube was then connected to the inlet-position of the 4-way Linear Connector (Dolomite microfluidics, UK connector (Part number 3000024). Droplets were pushed into the chip at 0.25 µl/min using a Legato 110 syringe pump (KD Scientific). During secondary droplet production, spacer oil was applied into the chip using a syringe system identical to that carrying the droplets, delivering oil to space the introduced droplets prior to the second emulsification. Spacer oil was connected to position 2 in the connector using a syringe pump set to deliver a flow of 0.40 µl/min. Double emulsion buffer (DE) was introduced to the chip using a Legato 100 (KD Scientific) single syringe pump applying pressure to a 10 mL syringe (Scientific Glass Engineering, Australia). Pump speed of the DE-buffer was set to 28 µl/min. Second emulsification was performed for 160 minutes until all primary droplets had passed the junction of the DE-chip.

### Droplet sorting and gating

Sorting was carried out on a single laser 488nm, S3e cell sorter (BioRad inc.) using ProSort software (v. 1.3b). Instrument PMTs were adjusted to: FSC=239, SSC=261, FL1=590, and FL2=367. FSC was used as primary threshold and the value was set to 1.00. Gating of positive droplets was performed in three consecutive gating events. First gate was set to discriminate between double emulsion droplets and “other” elements in the carrier buffer. The second gate was used to split the double emulsion droplets from the first gate into fluorescent and non-fluorescent double emulsion droplets. The third gate was applied to ensure that only positive droplets with the expected properties were sorted.

Sorting purity was set to “Enrich” and event rate was kept as close to 4000 events/second as possible throughout the experiment. Prior to sorting droplets, 5 µl Tris (10 mM) was placed at the bottom of the 1.5 mL collection tube to avoid disrupting the sorted droplets. Sorting was done for a period of 27 minutes and a total of 143 positive double emulsion droplets were sorted. Upon completed sorting, the collection tube was centrifuged at 1000 g for 10 seconds to collect any liquid from the side of the tube, arising from the splash impact of sorted droplets hitting the liquid surface of inside tube.

### DNA amplification

The collected droplets were coalesced by adding 20µl of PicoBreak (Sphere Fluidics), mixing and centrifuging the sample. 3µl of the resulting aqueous-phase was used as template for a multiple displacement amplification (MDA) reaction. The MDA reaction mix was kindly provided by Samplix. The MDA reaction was emulsified on a x-junction droplet generator chip (ChipShop) using 1% PicoSurf <u>(Sphere Fluidics)</u> in 7500-Novec oil as carrier phase. The droplet production was driven by air pressure controlled by pressure regulator (Fluigent). The droplets containing the MDA reaction were incubated for 16 hours at 30C followed by 10min at 65C to terminate the reaction. 6ul from the MDA reaction was used as template for a second droplet MDA reaction. After each MDA round the emulsions where coalesced using 20µl PicoBreak.

### PacBio sequencing

A 2 kb PacBio library was produced from 1 µg enriched DNA using the SMRTbell™ Template Prep Kit 1.0 according to manufacturer’s instructions. The library was sequenced on one SMRTcell™ on the PacBio RSII instrument using C4 chemistry and P6 polymerase and 240 minutes movie time.

### Illumina sequencing

The enriched sample was diluted 10-fold with Tris-HCl to reduce the MgCl2 concentration before library preparation. A standard DNA library for Illumina was generated by GATC and 5 million read pairs of 2x 150bp were generated.

### Nanopore sequencing

An ONT library was produced from 400 ng of enriched DNA using the Rapid Sequencing (SQK-RAD003) protocol. The library was sequenced on 1 MIN106 flowcell (R9.4) for 17.5 hours with subsequent Albacore basecalling (v2.1.3). All was performed using standard settings according to manufacturer’s recommendations.

### Detection of HPV18 integration sites

HPV18 integration sites were identified by mapping all sequence reads to the HPV18 reference genome. The reads mapping to HPV18 were subsequently re-mapped to the human reference genome GRCh38 (hg38). Reads mapping to both genomes were considered HPV18/Chr8 fusion-reads.

### Sanger sequencing

PCR primers located on each side of the fusion points were designed and used to generate PCR amplicons across the fusion point. These PCR products were Sanger sequenced using one of the PCR primers.

## References

1. Alkan, C., Coe, B.P. & Eichler, E.E. Genome structural variation discovery and genotyping. Nat Rev Genet 12, 363–76 (2011).

2. Jain, M. et al. Nanopore sequencing and assembly of a human genome with ultra-long reads. Nat Biotechnol 36, 338–345 (2018).

3. Seo, J.S. et al. De novo assembly and phasing of a Korean human genome. Nature 538, 243– 247 (2016).

4. Mamanova, L. et al. Target-enrichment strategies for next-generation sequencing. Nat Methods 7, 111–8 (2010).

5. Ardui, S. et al. Detecting AGG Interruptions in Male and Female FMR1 Premutation Carriers by Single-Molecule Sequencing. Hum Mutat 38, 324–331 (2017).

6. Lode, L. et al. Single-molecule DNA sequencing of acute myeloid leukemia and myelodysplastic syndromes with multiple TP53 alterations. Haematologica 103, e13–e16 (2018).

7. Wang, M. et al. PacBio-LITS: a large-insert targeted sequencing method for characterization of human disease-associated chromosomal structural variations. BMC Genomics 16, 214 (2015).

8. Gabrieli, T., Sharim, H., Michaeli, Y. & Ebenstein, Y. Cas9-Assisted Targeting of CHromosome segments (CATCH) for targeted nanopore sequencing and optical genome mapping. bioRxiv (2017). http://dx.doi.org/10.1101/110163

9. Tsai, Y.-C. et al. Amplification-free, CRISPR-Cas9 Targeted Enrichment and SMRT Sequencing of Repeat-Expansion Disease Causative Genomic Regions. bioRxiv (2017). http://dx.doi.org/10.1101/203919

10. Hoijer, I. et al. Detailed analysis of HTT repeat elements in human blood using targeted amplification-free long-read sequencing. Hum Mutat (2018). http://dx.doi.org/10.1002/humu.23580

11. Adey, A. et al. The haplotype-resolved genome and epigenome of the aneuploid HeLa cancer cell line. Nature 500, 207–11 (2013).

12. Schneider-Gadicke, A. & Schwarz, E. Different human cervical carcinoma cell lines show similar transcription patterns of human papillomavirus type 18 early genes. EMBO J 5, 2285–92 (1986).

13. Travers, K.J., Chin, C.S., Rank, D.R., Eid, J.S. & Turner, S.W. A flexible and efficient template format for circular consensus sequencing and SNP detection. Nucleic Acids Res 38, e159 (2010).

14. Huddleston, J. et al. Reconstructing complex regions of genomes using long-read sequencing technology. Genome Res 24, 688–96 (2014).

15. Albrecht, V. et al. Dual redundant sequencing strategy: Full-length gene characterisation of 1056 novel and confirmatory HLA alleles. HLA 90, 79–87 (2017).

16. Loomis, E.W. et al. Sequencing the unsequenceable: expanded CGG-repeat alleles of the fragile X gene. Genome Res 23, 121–8 (2013).

17. Ameur, A., Kloosterman, W.P. & Hestand, M.S. Single-Molecule Sequencing: Towards Clinical Applications. Trends Biotechnol (2018). doi: 10.1016/j.tibtech.2018.07.013

